# The impact of Spike mutations on SARS-CoV-2 neutralization

**DOI:** 10.1101/2021.01.15.426849

**Authors:** C Rees-Spear, L Muir, SA Griffith, J Heaney, Y Aldon, JL Snitselaar, P Thomas, C Graham, J Seow, N Lee, A Rosa, C Roustan, CF Houlihan, RW Sanders, R Gupta, P Cherepanov, H Stauss, E Nastouli, on behalf of the SAFER Investigators, KJ Doores, MJ van Gils, LE McCoy

## Abstract

Multiple SARS-CoV-2 vaccines have shown protective efficacy, which is most likely mediated by neutralizing antibodies recognizing the viral entry protein, Spike. Antibodies from SARS-CoV-2 infection neutralize the virus by focused targeting of Spike and there is limited serum cross-neutralization of the closely-related SARS-CoV. As new SARS-CoV-2 variants are rapidly emerging, exemplified by the B.1.1.7, 501Y.V2 and P.1 lineages, it is critical to understand if antibody responses induced by infection with the original SARS-CoV-2 virus or the current vaccines will remain effective against virus variants. In this study we evaluate neutralization of a series of mutated Spike pseudotypes including a B.1.1.7 Spike pseudotype. The analyses of a panel of Spike-specific monoclonal antibodies revealed that the neutralizing activity of some antibodies was dramatically reduced by Spike mutations. In contrast, polyclonal antibodies in the serum of patients infected in early 2020 remained active against most mutated Spike pseudotypes. The majority of serum samples were equally able to neutralize the B.1.1.7 Spike pseudotype, however potency was reduced in a small number of samples (3 of 36) by 5–10-fold. This work highlights that changes in the SARS-CoV-2 Spike can alter neutralization sensitivity and underlines the need for effective real-time monitoring of emerging mutations and their impact on vaccine efficacy.

## Introduction

Serum neutralization activity is a common correlate of protection against viral infection following vaccination or natural infection (Plotkin, 2008). However, effective protection from viral infection can also require sufficient breadth of serum neutralization rather than potency alone. This is because of the high-levels of variation observed in major antigens across some viral populations (Burton, Poignard, Stanfield, & Wilson, 2012). The classic example in which a lack of breadth limits the protective capacity of the antibody response is Influenza. Here, the majority of neutralizing serum antibodies target only a particular set of influenza strains as a result of antigenic drift of the immunodominant hemagglutinin head (Zost, Wu, Hensley, & Wilson, 2019). Due to this, an annual vaccine is required and must be matched to the most probable circulating strain in any given year to ensure protection from infection. Emerging data from human vaccine trials and challenge studies in animal models suggest that neutralizing antibodies can prevent disease caused by infection with SARS-CoV-2, the virus that causes COVID-19 (McMahan et al., 2020; Polack et al., 2020). However, new variants of SARS-CoV-2 have begun to emerge in both human and farmed animal populations (S. Kemp et al., 2020; Oude Munnink et al., 2020; Tegally et al., 2020; Welkers, Han, Reusken, & Eggink, 2020). These variants include mutations in the major neutralizing antigen, the Spike glycoprotein, and raises the question of whether neutralizing serum responses induced by early circulating strains or by vaccines based on the Spike sequence of these early strains can neutralize the recently emerged virus variants.

Prior to the emergence of multiple mutations in Spike in the human population, we reasoned that a logical way to identify potential escape mutations was to look at sites of amino acid variation relative to the most closely related human betacoronavirus SARS-CoV, which caused the original SARS outbreak in 2003 (CDC, 2003). These two closely related viruses are characterized by a notable difference in transmission dynamics and disease outcomes (Cevik, Kuppalli, Kindrachuk, & Peiris, 2020; Lipsitch et al., 2003; Petersen et al., 2020) but both use the human ACE2 protein as a viral entry receptor (W. Li et al., 2003) and share approximately 75% similarity overall in Spike at an amino acid level (Gralinski & Menachery, 2020). Both viruses use the same region of their respective Spikes to bind ACE2; the receptor binding domain (RBD - found within the S1 subunit of Spike). There is considerable amino acid variation between the two RBDs, despite their conserved binding to ACE2, which explains why the majority of SARS-CoV-induced neutralizing monoclonal antibodies (mAbs) were found not to neutralize SARS-CoV-2, although some cross-binding activity has been observed (Wu et al., 2020) and targeted mutations can enable neutralization (Liu et al., 2020). Similarly, the majority of COVID-19 sera have either weaker or no neutralizing activity against SARS-CoV, but cross-neutralizing mAbs have been isolated (Brouwer et al., 2020).

Since the start of the pandemic, sequencing of viral populations has been deployed to enable detection of individual mutations within SARS-CoV-2 and identify new variants or strains that become dominant. Most recently, a new variant, B.1.1.7, has emerged in the UK (S. Kemp et al., 2020; Rambaut et al., 2020) that includes multiple mutations in both the RBD and the N-terminal domain (NTD) of Spike, both targets for neutralizing antibodies. Similarly, two further variants have been identified in South Africa, 501Y.V2, and Brazil, P.1, which carry additional mutations in both the NTD and RBD (Faria et al., 2021; Tegally et al., 2020). All three variants share a deletion of 3 amino acids in Orf1ab and key mutations in the RBD (E484K and the N501Y); data so far consistent with convergent evolution. Early reports have indicated that while the RBD mutation N501Y in the B.1.1.7 strain does not compromise post vaccine serum neutralization (Xie et al., 2021), one of the individual changes in the 501Y.V2 strain does impair neutralization but does not remove all activity (Greaney et al., 2021). Moreover, *in vitro* escape studies have shown similar mutations occur under selective pressure (Andreano et al., 2020).

Therefore, in this study we evaluated the potential role of individual amino acids in facilitating escape from neutralizing antibodies. Firstly, by making a series of point mutations to change the amino acids in SARS-CoV-2 to those found at the analogous position in SARS-CoV. Secondly, by making individual point mutations emerging in real world populations and by generating a pseudotype virus using the B.1.1.7 variant Spike sequence. We identify multiple mutations that can abrogate neutralization by some monoclonal antibodies targeting the RBD of Spike. However, in contrast, we show that serum responses are more resilient to these mutations, especially following severe infection where the antibody response is characterized by increased breadth.

## Results

### Generation of potential escape mutants by SARS-CoV amino acids substitution

There are 56 individual amino acid changes between the RBD of SARS-CoV-2 and SARS-CoV (Ortega, Serrano, Pujol, & Rangel, 2020), including sites at which antibody escape has been observed for SARS-CoV (Rockx et al., 2010). We identified 15 sites where single, or sequential, non-conservative amino acid changes were observed compared to SARS-CoV. These sites were mutated in the SARS-CoV-2 Spike to match SARS-CoV (Fig S1). Mutated Spike protein plasmids were then co-transfected with a lentiviral construct encoding luciferase and a packaging plasmid to produce pseudotyped viruses (Seow et al., 2020). Three of the substitutions resulted in virus that did not give sufficient titer to evaluate the impact on neutralization activity. The remaining mutated pseudotypes were then screened for any alteration in neutralization against a panel of human mAbs (Brouwer et al., 2020) isolated post SARS-CoV-2 infection. These mAbs have been previously mapped into eleven binding clusters whereby mAbs within a cluster reciprocally compete for binding to Spike. Of these, eight clusters comprise neutralizing mAbs (I, III, IV, VI, VII, IX, X, XI), five of which target the RBD (I, III, VI, VII, IX), and three clusters contain only non-neutralizing mAbs (II, V and VIII). Representatives of each neutralizing cluster were selected for evaluation against the Spike mutant pseudotypes.

### Impact of SARS-CoV Spike substitutions on SARS-CoV-2 mAb neutralization

Initial screening assays of the twelve infectious viral pseudotype mutants showed no effect on neutralization by any of the mAbs against three mutants with changes at amino acid positions RFA_346-8_KFP, S_459_G, and ST_477-8_GK (FigS1). In contrast, the remaining nine viral pseudotype mutants diminished neutralization for at least one mAb (Fig1) as described below:

**Fig 1.**
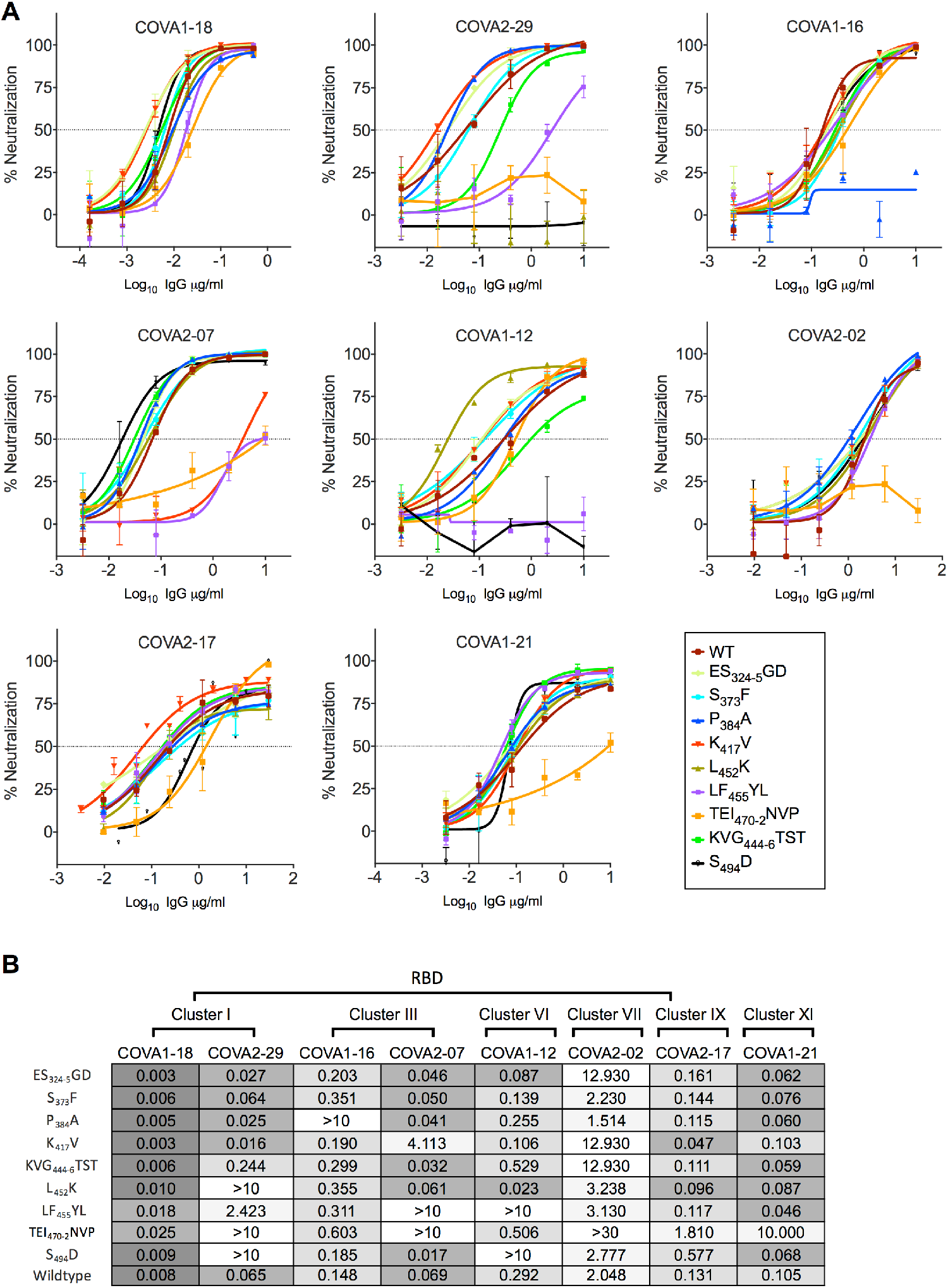
Mutating amino acids in SARS-CoV-2 Spike to match SARS-CoV decreases mAb neutralization. **(A)** Indicated mAbs were serially diluted in duplicate and incubated with each mutant SARS-CoV-2 luciferase-encoding pseudotyped virus (as noted in the legend) prior to the addition of HeLa cells expressing ACE-2. After two days, neutralization was measured as the relative reduction in relative light units (RLU). Data are representative of three independent repeats. The horizontal dotted line on each graph indicates 50% neutralization. **(B)** 50% inhibitory concentration (IC_50_) values were calculated for each mAb against the mutant SARS-CoV-2 pseudotyped viruses indicated in the left hand column. IC_50_ values are color-coded as follows: pale grey >1μg/ml, light grey 0.1-1 μg/ml, medium grey 0.1-0.01 μg/ml and dark grey <0.01 μg/ml. The previously established binding cluster for each mAb is indicated above each column, and whether or not the mAb binds RBD is also indicated.

### P_384_A

The P_384_A substitution resulted in complete loss of neutralization by COVA1-16 (Fig1A), a cluster III RBD-specific mAb that allosterically competes with ACE2 rather than directly blocking the binding site (Liu et al., 2020). Of note, this mutation has been described and structurally characterized elsewhere (Wu et al., 2020), revealing that this proline to alanine change results in a relatively small alteration in protein structure that can enable SARS-CoV mAbs to neutralize SARS-CoV-2 P_384_A. However, P_384_A does not weaken neutralization by any other mAbs, including another mAb in the cluster III competition group.

### K_417_V

The K_417_V mutation results in a pseudotyped virus that is less susceptible to COVA2-07 mediated neutralization, which is the other cluster III RBD-specific mAb screened (Fig1A). COVA2-07 belongs to the same sub-cluster as COVA2-04, the structure of which has been solved and contact residues include numerous bonded and non-bonded contacts within the RBD and ACE2-binding site (Wu et al., 2020). That this mutation should alter mAbs such as COVA2-07, which competes directly with ACE2 for binding, is not unexpected as the lysine at position 417 forms a hydrogen bond with ACE2 (Lan et al., 2020) that is likely disrupted by this substitution. We then evaluated an additional mAb, COVA2-04, from the same competitive binding cluster as COVA2-07. This is because COVA2-04 is representative of class of SARS-CoV-2 neutralizing antibodies that all use the VH3-53 gene and have been suggested to be an unusual “public” or stereotyped antibody (Cao et al., 2020; Mor et al., 2020; Robbiani et al., 2020). COVA2-04 was not able to neutralize the K_417_V pseudotype (data not shown). Although, of note, this mutation has no real impact on neutralization by any of the other mAbs tested.

### KVG_444-6_TST

This multiple substitution, which is a substantial change between SARS-CoV-2 and SARS-CoV, results in a 3.7-fold drop in neutralization potency for COVA2-29, which is a cluster I RBD-specific antibody (Fig1A, B). This is the largest effect of this mutation, as the neutralization activity of the other mAbs is largely unaffected despite the alteration of three sequential amino acids. This may be explained by the relatively minor differences in the amino acid side chains at the mutated residues. Unlike some of the other mutants tested, such as LF_455_YL that introduces a phenol, the substitutions made here may not disrupt protein structure or antibody binding to any great extent. Notably, binding analysis has suggested mutations in this loop can reduce RBD binding by Spike-specific sera (Greaney et al., 2021).

### L_452_K

This mutation is situated directly within the receptor binding motif (RBM) of the RBD. It renders pseudotyped virus resistant to neutralization by the cluster I mAb COVA2-29, but does not affect the other cluster I mAb COVA1-18 or any other mAb tested.

### LF_455_YL

This double substitution reduces neutralization by RBD-specific mAbs from different clusters, specifically, the cluster I mAb COVA2-29, cluster III mAb COVA2-07 and cluster VI mAb COVA1-12 (Fig1A, B). For COVA1-12 all neutralization activity is abolished, while COVA2-07 activity is just below the level required to calculate an IC_50_ value.

### TEI_470-2_NVP

This triple mutation is located in a loop within the RBM where other substitutions have been reported to abolish ACE2 binding (Xu et al., 2021; Yi et al., 2020). This mutation prevents neutralization by COVA2-29 (cluster I), COVA2-07 (cluster III) and COVA2-02 (cluster VII). It also reduces the activity of the most potent mAb COVA1-18 (cluster I) by 3-fold, whereas this mAb is only minimally affected by other mutations. Moreover, TEI_470-2_NVP lowers the potency of the structurally unmapped non-RBD cluster XI mAb, COVA1-21, to the limit of detection. Thus, this mutation negatively impacts the most mAbs, including representatives from four separate epitope clusters.

### S_494_D

This single substitution towards the end of the RBM destroys neutralization activity by both COVA2-29 (cluster I) and COVA1-12 (cluster VI) but does not have a major impact on the other cluster I mAb tested or those from other epitope clusters.

In summary, different mAbs can lose their neutralization activity when confronted with different Spike mutations and the effects are not strictly delineated by binding clusters, such that mAbs within the same competition cluster are frequently differentially affected. The triple substitution TEI_470-2_NVP has the most detrimental effects on different antibodies and impacts mAbs from nearly all binding clusters. LF_455_YL also negatively affects mAbs across different binding clusters. Notably, COVA2-29 is the mAb that is most frequently negatively affected by different mutations, with substantial loss of some neutralizing activity against four mutations (L_452_K, LF_455_YL, TEI_470-2_NVP, S_494_D), which are all within the RBM of the RBD (Fig1A). COVA2-29 is also known to compete directly with ACE2 for binding to Spike, which may explain why it is sensitive to so many different mutations within the RBM.

### Impact of Spike mutations on serum neutralization

Following the identification of Spike mutations that can limit or abrogate neutralizing activity of mAbs (Fig2A), the next step was to assess the impact of these mutations on serum neutralization. Samples were tested following two different infection scenarios. Firstly, from a previously characterized cohort of seropositive healthcare workers who experienced mild or asymptomatic SARS-CoV-2 infection (Houlihan et al., 2020). Secondly, sera were obtained from a cohort of hospitalized patients who experienced severe disease. Eighteen samples were chosen from both cohorts for screening purposes to obtain representatives with intermediate (1:50-100), strong (1:100-1000) and potent (>1:1000) neutralizing ID_50_ values. The median serum ID_50_ for hospitalized patients selected was 1:1275, and that for selected mild/asymptomatic cases was 1:1045. Strikingly, serum samples from both cohorts are less impacted by Spike mutations than individual mAbs in terms of fold decrease in neutralization potency (Fig2B and C). Only one of thirty-six serum samples lost all neutralizing activity (Fig S2), in contrast to the five mAbs from five different epitope clusters where neutralization was completely abrogated by a single Spike mutation (Fig 1B). Moreover, fold-decrease in neutralization potency was more modest for sera than mAbs, with an average 2-fold decrease across all sera for the most disadvantageous mutation TEI_470-2_NVP as compared to a more than 100-fold decrease observed for several of the mAbs (Fig2B and C). Interestingly, none of the 36 serum samples lost >5-fold potency against the other triple substitution, KVG_444-6_TST which contrasts with recent data showing a single mutation at G_446_ caused a major loss of neutralization in one sample (Greaney et al., 2021). Importantly, there was notable difference between the resilience of serum samples from severely ill hospitalised individuals and those who had experienced mild/asymptomatic infection. Only two serum sample from a hospitalised individual lost more than 3-fold potency against any individual mutant (Fig2C, FigS2). Whereas approximately half (10 of 18) the mild/asymptomatic serum samples showed a three-fold drop in potency against at least one Spike mutant (Fig2C, Fig S2).

**Fig 2.**
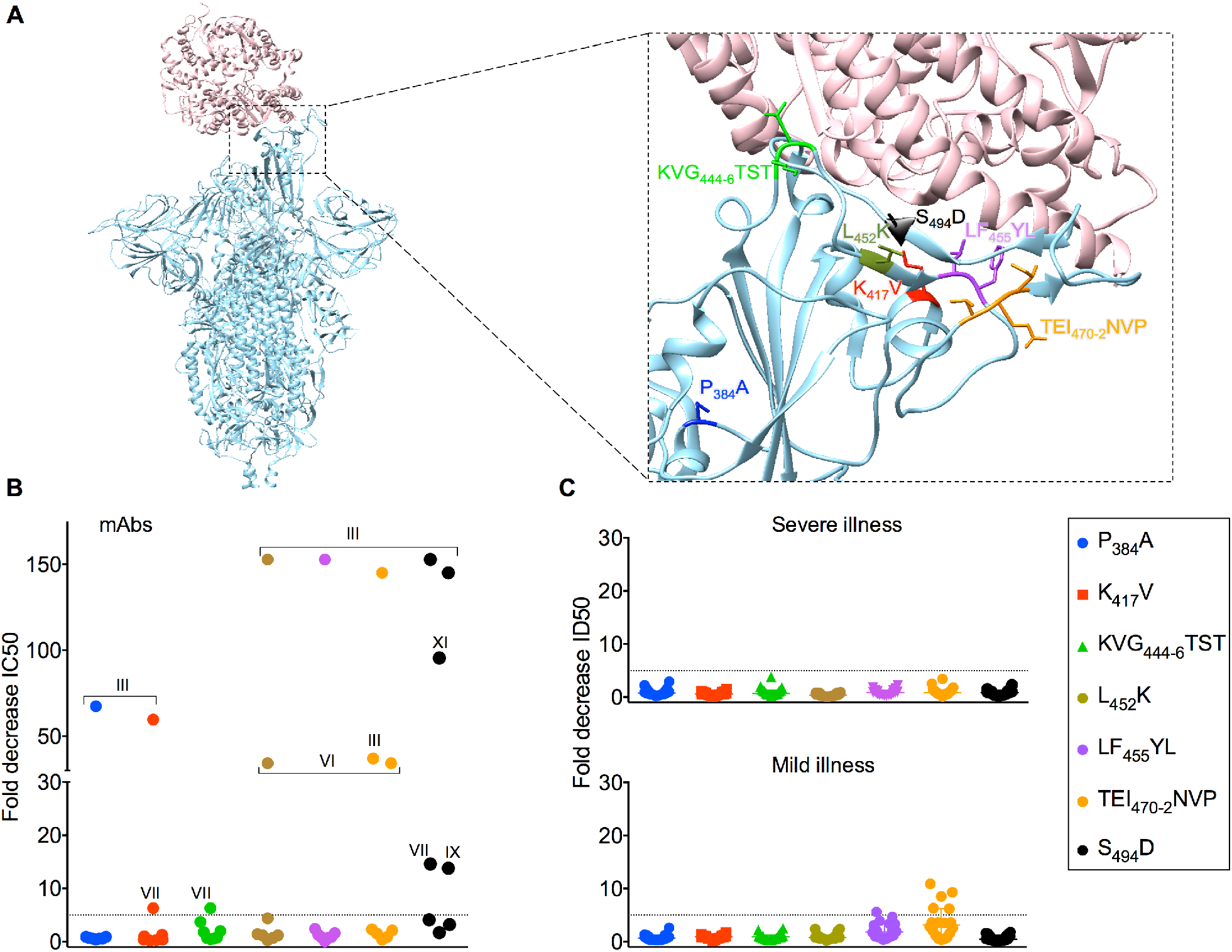
Neutralization by sera is less adversely affected by SARS-CoV amino acid substitutions in SARS-CoV-2 Spike. **(A)** Representation of SARS CoV-2 Spike trimer (blue) in complex with ACE-2 (pink) (PDB code 7DF4). Magnified image shows mutated amino acid side chains at residues of interest. **(B)** IC_50_ values for each mAb against SARS-CoV-2 wildtype pseudotyped virus were divided by the IC_50_ for each mutant pseudotyped virus against the corresponding mAb to generate the fold decrease in neutralization on the Y-axis. The dotted horizontal line indicates a 5-fold drop in neutralization potency. The competitive binding clusters of each mAb that loses > 5-fold neutralization activity are labeled on the graph. **(C)** Thirty six serum samples were serially titrated and incubated with the mutant SARS-CoV-2 luciferase-encoding pseudotyped viruses indicated in the legend prior to the addition of HeLa cells expressing ACE-2. After two days, neutralization was measured as the relative reduction in relative light units (RLU) and 50% inhibitory dilution factors calculated using Graphpad Prism. ID_50_ values for each sera against SARS-CoV-2 wildtype pseudotyped virus were divided by the ID_50_ for each mutant pseudotyped virus against the corresponding sera to generate the fold decrease in neutralization on the Y-axis. The dotted horizontal line indicates a 5-fold drop in neutralization potency. The 18 serum samples from hospitalized patients are shown in the upper graph labeled “severe illness” and the 18 serum samples from healthcare workers who experience mild/asymptomatic COVID-19 are shown in the lower graph labeled “mild illness”.

### Greater levels of Spike-reactive antibodies in sera after severe illness

The differences in resilience to Spike mutations seen in the neutralizing sera from these two infection scenarios is plausibly due to greater polyclonality arising from greater antigenic stimulation during severe illness. To assess the serological profiles of these two cohorts, we compared the 50% inhibitory dilution (ID_50_) values across 192 samples and measured the binding titers by semi-quantitative ELISA for 199 samples as previously described (Ng et al., 2020; O’Nions et al., 2020). There is a significantly higher median IgG binding titer of 46.5 μg/ml following severe illness versus 3.9 μg/ml following mild/asymptomatic disease (Fig3A, FigS3). Similarly, there is a significantly higher median ID_50_ value in the hospitalized patient cohort as compared to the mild/asymptomatic group (Fig3B). This shows, as has been observed previously (Seow et al., 2020), that severely ill patients have higher binding and neutralization titers than asymptomatic/mild cases and that higher binding titers correlate with greater neutralization. However, when considering how the IgG binding titer from each individual relates to their neutralization titer it became clear that there was a discrepancy (Fig3C and D). Most hospitalized patients required a binding titer of greater than 10 μg/ml to achieve strong neutralization (ID_50_ >100). Moreover, mild infections could lead to potent neutralization (ID_50_ >1000) at binding titers of less than 10 μg/ml (Fig3D) whereas this was observed for only two individuals following severe illness (Fig3C). In fact, the amount of specific IgG present at the serum ID_50_ is significantly higher in severe illness compared to mild disease (Fig 3E). This suggests that a higher proportion of antibodies are non-neutralizing following severe illness. However, the total level of specific antibodies is so high that the number of neutralizing antibodies following severe illness may be greater and explain the relative resilience of severe sera to Spike mutations.

**Fig 3.**
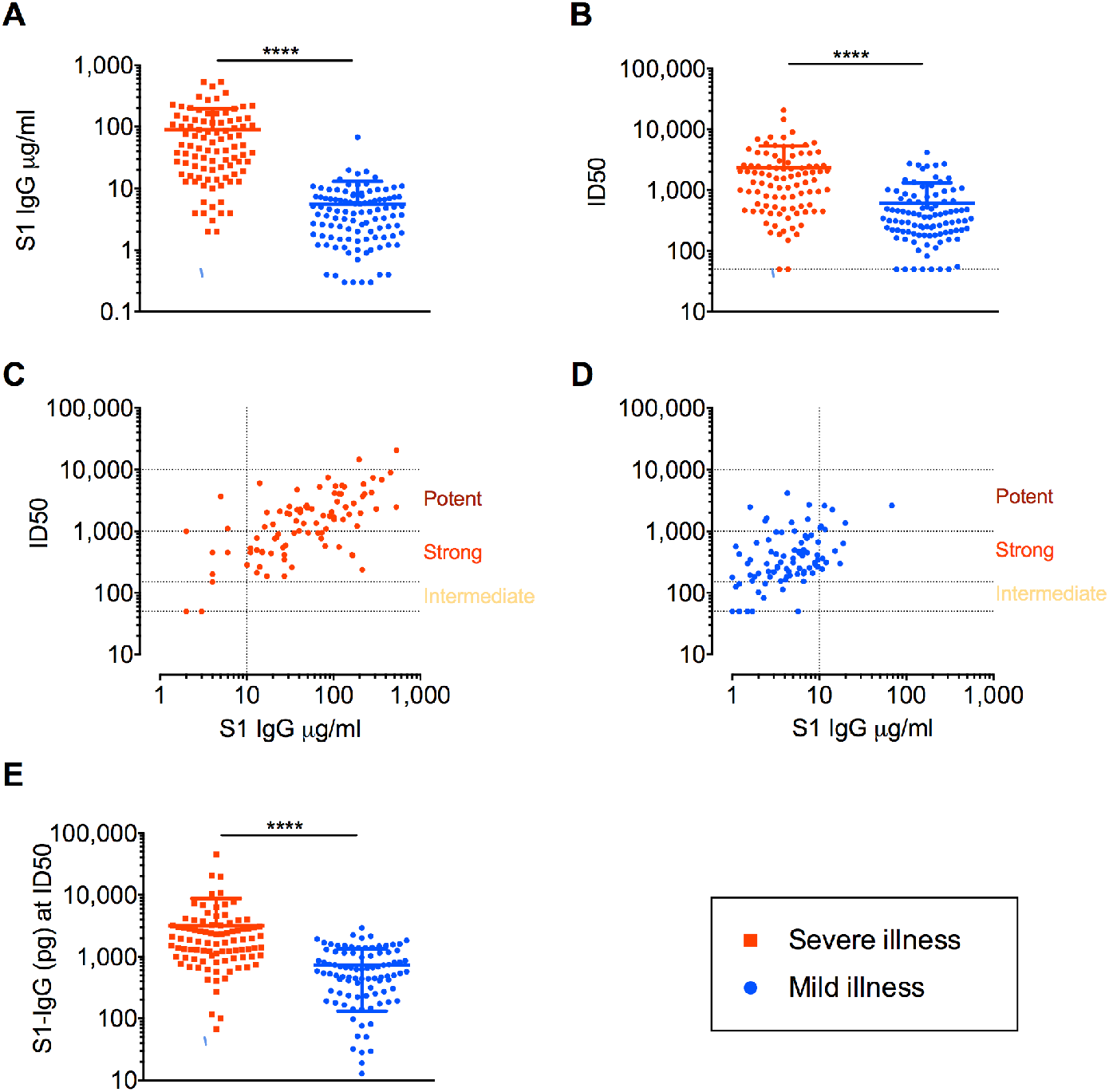
Serum responses following severe COVID-19 have greater polyclonality but less efficient neutralization. **(A)** Spike S1 subunit semi-quantitative titers measured by ELISA (see Methods) are shown on the Y-axis for 94 serum samples from hospitalized COVID-19 patients and 105 serum samples from healthcare workers who experienced mild COVID-19 disease. **(B)** Serum ID_50_ values measured by pseudotyped neutralization assay (see Methods) are shown on the X-axis for 93 serum samples from hospitalized COVID-19 patients and 99 serum samples from healthcare workers who experienced mild COVID-19 disease. Note, some serum samples from the original cohort that had binding titers gave no neutralization titer (6 from healthcare workers, 1 from a hospitalized patient). **(C)** ID_50_ values measured by pseudotyped neutralization assay for serum samples from hospitalized COVID-19 patients plotted on the Y-axis against the corresponding S1 IgG binding titer for each sample. Relative ranking of neutralization titers is indicated on the graph. (**D)** Serum ID_50_ values measured by pseudotyped neutralization assay for the serum samples from healthcare workers who experienced mild COVID-19 disease are plotted on the Y-axis against the corresponding S1 IgG binding titer for each sample. Relative ranking of neutralization titers is indicated on the graph. Only sera that gave a measurable titer in both semi-quantitative ELISA or pseudotype neutralization assay were included in (B), (C) and (D). Serum sample groups are color-coded according to the legend. **(E)** Concentrations of S1-specific serum IgG (pg) at ID_50_ dilutions were calculated using the IgG titers quantified via the semi-quantitative ELISA and the known ID_50_ value. Only sera that gave a measurable titer in both semi-quantitative ELISA and pseudotype neutralization assay were included. Data for (A)–(B) and (E) were analyzed by a non-parametric Mann-Whitney U test.

### Impact of Spike variants on mAb and serum neutralization

Investigating the ability of post-SARS-CoV-2-infection mAbs and serum to cope with mutations in Spike engineered based on differences with SARS-CoV was a rational first approach to study escape, because these mutations were likely to form viable Spike proteins. Indeed, some of the positions at which amino acids were mutated, as described above, have now been observed to be a site of variance in SARS-CoV-2 across the human population (Q. Li et al., 2020; Liu et al., 2020; Starr et al., 2020). However, additional viral variants, not linked to points of variance between these two closely related viruses, have started to emerge on a significant scale (Q. Li et al., 2020; Weisblum et al., 2020). Firstly, the D_614_G mutation, observed in western Europe in February 2020 and now dominant across the globe (Korber et al., 2020). It has already been described that D_614_G has higher infectivity and greater viral replication (Korber et al., 2020; Q. Li et al., 2020; Plante et al., 2020) and somewhat increases the ability of serum and mAbs to neutralize SARS-CoV-2 (Korber et al., 2020; Plante et al., 2020; Weissman et al., 2020). More recently, a new variant of SARS-CoV-2 (B.1.1.7) has emerged in England and been associated with a rapid rise in case numbers (S. Kemp et al., 2020; Rambaut et al., 2020) and high viral loads (Kidd et al., 2020). B.1.1.7 encodes 8 sites of change in Spike relative to the original Wuhan strain. Of these, the most likely candidates to alter neutralization sensitivity are the deletion in the NTD (ΔH_69_/V_70_) and the N_501_Y substitution in the RBM (S. Kemp et al., 2020; Rambaut et al., 2020). Therefore, these changes were introduced in the Wuhan strain Spike in the presence of D_614_G to produce pseudotyped viruses for neutralization sensitivity analysis. Firstly, ΔH_69_/V_70_ did not negatively impact the neutralization potency for most of the mAbs tested, including cluster IX mAb COVA2-17 (Fig4A), which has recently been found to bind the NTD (Rosa et al., 2021). The exception was the structurally unmapped COVA1-21 (cluster XI), which was previously reported to lose partial potency against this deletion in the context of other mutations that arose in an immuno-compromised patient (S. A. Kemp et al., 2020). Similarly, the serum neutralization activity did not decrease by more than 3-fold relative to the D_614_G Spike for any individual sample (Fig4B). In contrast, the introduction of the N_501_Y substitution (observed in both B.1.1.7, 501Y.V2 and P.1) lowered the neutralization potency of one mAb, COVA1-12, to the limit of the assay, with a fold decrease in IC_50_ of >40 (Fig4A). This cluster VI RBD-specific mAb was also negatively impacted by proximal mutations at LF_455_YL and S_494_D. Moreover, a 5-fold decrease in potency was observed against the N_501_Y pseudotype for the cluster IX mAb COVA2-17, which also loses efficacy against TEI_470-2_NVP. However, as seen for the other mutations that abrogate mAb function, the N_501_Y change had less of an effect on the sera obtained after both severe and mild infection, with all individual serum samples remaining within three-fold of their original ID_50_ value (Fig4B, FigS4).

**Fig 4.**
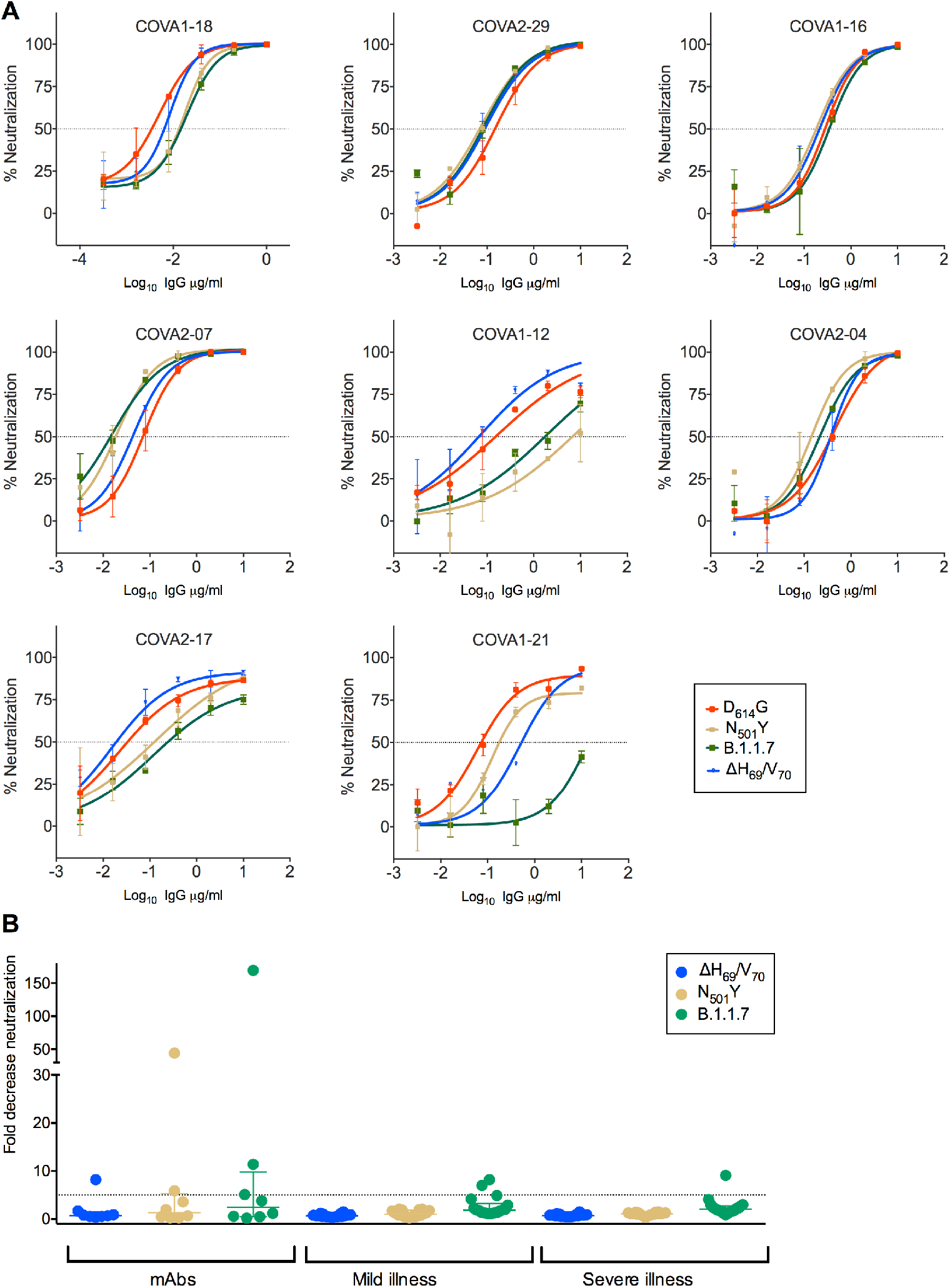
Variant B.1.1.7 SARS-CoV-2 Spike effect on mAb and serum neutralization. **(A)** Indicated mAbs were serially diluted in duplicate and incubated with the mutant SARS-CoV-2 luciferase-encoding pseudotyped virus in the legend prior to the addition of HeLa cells expressing ACE-2. After two days neutralization was measured as the relative reduction in relative light units (RLU). Data are representative of three independent repeats. The horizontal dotted line on each graph indicates 50% neutralization. **(B)** IC_50_ values for each mAb or ID_50_ values for each serum sample against SARS-CoV-2 D_614_G pseudotyped virus were divided by the IC_50_ for each mutant pseudotyped virus against the corresponding mAb to generate the fold decrease in neutralization on the Y-axis, as color-coded in the key. The dotted horizontal line indicates a 5-fold drop in neutralization potency. Whether fold decrease in neutralization potency refers to mAbs, 18 serum samples from hospitalized patients or 18 serum samples from healthcare workers who experience mild/asymptomatic COVID-19 is indicated under the graph by the labels “mAbs”, “severe illness” and “mild illness”, respectively.

### Impact of B.1.1.7 Spike on mAb and serum neutralization

Finally, a B.1.1.7 Spike pseudotyping plasmid was synthesized to incorporate the mutations observed in this new variant in combination, and then evaluate neutralization sensitivity of mAbs and sera. This showed that, as per individual mutants, B.1.1.7 can lessen the potency of some mAbs, although unlike other mutations described above, it does not remove all activity for any individual mAb and in total three mAbs were affected, COVA2-17, COVA1-12 and COVA1-21 (Fig4A). These mAbs belong to distinct clusters and so do not compete for binding to the same epitope. Firstly, the cluster IX mAb COVA2-17 showed an approximate 5-fold drop in potency against both the N_501_Y single mutant and the B.1.1.7 pseudotype, implying this loss of potency is primarily N_501_Y driven. In contrast, the decrease in potency noted with the single N_501_Y change for the RBD-specific mAb COVA1-12 was lessened in the context of B.1.1.7, with a 11-fold rather than a >40-fold drop in neutralization. Furthermore, one mAb, COVA1-21 experienced a substantial drop in potency against B.1.1.7 compared to the single mutants or the D_614_G spike. This cluster XI mAb, which does not bind to RBD or S1 subunits, showed more than 100-fold reduction in potency. This effect is likely in part mediated by the ΔH_69_/V_70_ deletion that also reduced the potency of COVA1-21 when evaluated as a single mutant. The B.1.1.7 Spike was then tested against the 36 serum samples from the two cohorts (Fig4B). The maximum fold-decrease in potency for the serum samples from mild illness was 8.2 but the majority of samples showed less than a 3-fold change. Similarly, the maximum decrease seen for samples from hospitalized patients was a 9.1-fold change, but most of the samples showed minimal change in the potency of their neutralization. Overall, three samples from each cohort (8%) showed a 5–10-fold reduction, but as they were potently neutralizing sera, the reduced ID_50_ values were still >1:100. Similarly, another four samples from both cohorts (11%) showed a 3–5-fold reduction in neutralization when tested against the B.1.1.7 Spike pseudotype. Again, the reduced ID_50_ values were still potent (on average >1:500) with only two samples having an ID_50_ of <1:200 (Fig S4)

## Discussion

This study demonstrates that Spike mutations can diminish or abolish neutralizing activity by individual mAbs but that serum neutralization is less strongly affected. Notably, only one engineered mutation, and none of the observed Spike mutants or the B.1.1.7 variant, resulted in a complete escape from neutralizing activity and this was only seen for one out of thirty-six serum samples. The Spike mutants evaluated comprise seven substitutions designed to mimic possible escape changes based on homology with SARS-CoV, two observed high-frequency mutations and the B.1.1.7 Spike variant. The most likely explanation for the greater effect on mAbs as compared to sera is the inherent polyclonality underlying serum neutralization. This concept is supported by the observation that single Spike mutations can weaken neutralization for a particular mAb but not for other mAbs from within the same binding cluster. This highlights that different antibodies use distinct molecular contacts within shared epitopes, such that a single mutation may not be detrimental to all antibodies within the same binding cluster. Thus, because polyclonal sera contain multiple antibodies that target the major neutralizing sites in subtly different ways, it is less sensitive to Spike mutations.

The Spike mutations studied here were designed to identify potential escape variants by mimicking in part the natural variation observed between SARS-CoV and SARS-CoV-2, and are focused mainly on the RBD as the major site of neutralizing antibody activity. Therefore, it was unsurprising that many of the RBD-specific mAbs evaluated here lost neutralization activity against one or more of these mutations. These included substantial multi-residue substitutions not yet seen in the SARS-CoV-2 but also single point mutations at positions that have been observed to mutate in the real world (S. Kemp et al., 2020; Korber et al., 2020; Rambaut et al., 2020; Tegally et al., 2020). These include, K_417_V that had a negative effect on mAb neutralization and another mutation at this position, K_417_N, has been observed in the emerging South African variant (Tegally et al., 2020). Importantly, the mAb (COVA2-07) that loses >40-fold neutralization activity against the K_417_V pseudotyped virus belongs to the same cluster as COVA2-04 that loses all neutralization activity against the K_417_V mutant. COVA2-04 belongs to the VH3-53 “public” BCR against SARS-CoV-2 identified from multiple human infections. Thus, COVA2-04-like antibodies are thought to be widespread among the seropositive population and so the K_417_V would be predicted to have an impact on sera from many individuals. Interestingly, serum samples from mild infection showed very little change in neutralization potency against this mutation. However, the strongest effect on serum samples from mild infection was mediated by the TEI_470-2_NVP substitution, which is part of what has been termed the RBD binding ridge, and other mutations in this region can decrease serum neutralization (Greaney et al., 2021). As such, any mutation in this zone should be closely monitored in viral populations due to the potential for escape. However, it is encouraging that the effect of these mutations on sera was much less pronounced and that individual mAbs within the same clusters were not universally inactivated by any given mutation. Notably, the mutations that most substantially decrease serum neutralization are those that negatively impact mAb activity against the widest range of clusters (I, III, XI, IX and VI) suggesting that mAb screening is a useful proxy for potential serum effects if a range of antibody clones are used. However, the capacity to predict the *in vivo* impact of a drop in neutralization potency requires correlation of *in vitro* serum neutralization ID_50_ values with protection, which thus far has only been achieved in animal models where, encouragingly, an ID_50_ value of 1:50 was found to be protective (McMahan et al., 2020).

A caveat to the first part of this study is that only RBD substitutions were considered. Further studies to assess potential mutations before they arise should include those in NTD given the emerging importance of NTD as a site for neutralizing antibodies (Andreano et al., 2020; Rosa et al., 2021). However, it should be noted that one RBD change, TEI_470-2_NVP, resulted in a 24-fold drop in potency for COVA1-21, which does not bind RBD and remains structurally unmapped (Brouwer et al., 2020). Regardless, that sera likely containing a mixture of RBD-and NTD-specificities are more resilient than individual mAbs in the face of Spike mutations within the RBD is not surprising. This further highlights the importance of a broad polyclonal serum response to maintain neutralizing activity in the event of novel Spike mutations emerging, and the need to consider more than RBD binding in serological evaluations.

To understand if the conclusions from studying the impact of the SARS-CoV-2/SARS-CoV substitutions on neutralization parallel those of real-world Spike mutations, we examined the responses to the newly emerged B.1.1.7 variant (S. Kemp et al., 2020; Rambaut et al., 2020). This revealed that the first NTD deletion observed, ΔH_69_/V_70_, did not alter RBD-specific mAbs or any sera. Although as previously described (S. A. Kemp et al., 2020) it did result in a drop in potency for non-RBD mAb COVA1-21. The RBD mutation N_501_Y, shared between B.1.1.7, 501Y.V2 and P.1, did remove almost all neutralizing activity for one mAb but, in a similar pattern to other substitutions, this did not translate to any large effect on serum potency. Of note, we have not studied changes at position 484 that have been observed in 501Y.V2 and P.1 and reported to reduce neutralization in serum samples (Greaney et al., 2021) and during *in vitro* escape (Andreano et al., 2020). Further studies of mutations at position 484 and new emerging mutations will be needed. This would best be facilitated by large, curated panels of mAbs and pools of sera from individuals with different infection/vaccination trajectories.

Theoretically it is likely that combinations of mutations have more potential to lead to loss of serum activity than individual single amino acid changes by destroying multiple parts of key epitopes. This has partially been observed in terms of the new B.1.1.7 Spike pseudotype analyzed here. Only one mAb was more dramatically affected by the full set of B.1.1.7 mutations. However, the small number of serum samples with reduced neutralization relative to the D_614_G virus, were more strongly affected by B.1.1.7 than either the ΔH_69_/V_70_ or N_501_Y mutations individually (Fig 4B). This reduced neutralization was seen in 11% (3–5-fold) and 8% (5–10-fold) when tested against the B.1.1.7 Spike pseudotype. However, all of the affected samples were still able to neutralize B.1.1.7, and the average reduced serum ID_50_ value was 1:522. This is ten-times higher than the reported serum ID_50_ correlate of protection in animal studies and suggests these responses would likely still be effective against infection with B.1.1.7.

The differences in the data observed with B.1.1.7 and the two individual mutations ΔH_69_/V_70_ or N_501_Y highlight the importance of testing emerging variants in the full form. Moreover, this approach may be important as combinations of mutations could enable individual antibody escape mutations that are disadvantageous for transmission to be propagated. For example, a residue such as S_494_ is involved both in ACE2 binding (Xu et al., 2021) and mAb neutralization (Fig1). Therefore, a mutation at S_494_ could decrease antibody function but also decrease host receptor recognition and limit infectivity. However, the detrimental effects of the mutation on infectivity could be compensated for if a S_494_ mutation occurred in concert with a mutation that strengthened a different part of the viral entry pathway as has been suggested(S. A. Kemp et al., 2020). This highlights the need for rapid evaluation of variant strains upon emergence, potentially accelerated by computational modeling based on prior knowledge of the effects of individual changes.

In conclusion, this study underlines both the potential for escape from neutralizing antibodies due to mutations in Spike and the relative resilience of serum responses compared to individual mAbs. This difference likely derives from the breadth inherent in polyclonal sera as compared to the precision interaction of a given mAb. Our results suggest that the majority of vaccine responses should be effective against the B.1.1.7 variant as the sera evaluated were obtained after infection early in the pandemic when the commonly circulating virus was highly similar in sequence to the vaccines now being deployed. A reduction in potency was observed in a minority of samples tested against B.1.1.7, however, neutralization titers remained above 1:200 in almost all cases. Finally, it is probable that as SARS-CoV-2 seropositivity increases across the human population (due to both vaccination efforts and natural infection) there could eventually be selection for Spike mutations that result in substantial antigenic drift as seen for Influenza. If and when this will happen is unpredictable given the current scale of ongoing transmission worldwide. The data herein suggest evaluation of neutralizing mAbs from non-overlapping binding clusters can highlight which Spike mutations will most impact sera neutralization. Greater knowledge of the molecular epitopes recognized by individual mAbs and their relative immunodominance within sera is needed urgently. This is because understanding of rules of engagement for SARS-CoV-2 neutralizing antibodies is a crucial component of preparedness for major antigenic changes and long-term management of coronaviruses globally. Our findings stress the importance of continuous monitoring of variants and in vitro assessment of their impact on neutralization. This is particularly relevant for the use of convalescent plasma and the development of therapeutic monoclonal antibodies as well as vaccine development and implementation.

## Acknowledgements

The authors would like to thank James E Voss for the gift of HelaACE2 expressing cells, George Kassiotis, Dan Frampton, Ann-Kathrin Reuschl and Joe Grove for helpful discussion and critical feedback. We are indebted to the Biobank staff and study participants and their families at the Royal Free Hospital, and the UCLH SAFER study recruitment team and study participants.

## Author contributions

KJD, LEM, LM, SAG, PT, NL and CR-S characterized monoclonal antibodies and sera; AR, CR, and PC expressed and purified proteins; JH, CH, HS and EN assembled the panels of human sera samples; MJvG, RWS, YA, JLS isolated and provided monoclonal antibodies; RG generated and provided mutated Spike plasmids, MJvG, KJD, and LEM wrote the paper with contributions from all authors.

## Methods

### Spike mutant generation

QuikChange Lightening Site-Directed Mutagenesis kit was used to generate amino acid substitutions in the SARS-CoV-2 Wuhan Spike expression vector (Seow et al., 2020) or the D614G pCDNA Spike plasmid (S. A. Kemp et al., 2020) following the manufacturer’s instructions (Agilent Technologies, Inc., Santa Clara, CA). Spike B.1.1.7 was synthesised by Genewiz, Inc. and cloned into the pCDNA expression vector using BamHI and EcoRI restriction sites.

### Neutralization assay

HIV-1 particles pseudotyped with SARS-Cov-2 spike were produced in a T75 flask seeded the day before with 3 million HEK293T cells in 10 ml complete DMEM, supplemented with 10% FBS, 100 IU/ml penicillin and 100 μg/ml streptomycin. Cells were transfected using 60 μg of PEI-Max (Polysciences) with a mix of three plasmids: 9.1 μg HIV-1 luciferase reporter vector (Seow et al., 2020), 9.1 μg HIV p8.91 packaging construct and 1.4 μg WT SARS-CoV-2 spike expression vector (Seow et al., 2020). Supernatants containing pseudotyped virions were harvested 48 h post-transfection, filtered through a 0.45-μm filter and stored at −80°C. Neutralization assays were conducted by serial dilution of monoclonal IgG at the indicated concentrations in DMEM (10% FBS and 1% penicillin–streptomycin) and incubated with pseudotyped virus for 1 h at 37°C in 96-well plates. HeLa cells stably expressing ACE-2 (provided by J.E. Voss, Scripps Institute) were then added to the assay (10,000 cells per 100 µl per well). After 48-72 h luminescence was assessed as a proxy of infection by lysing cells with the Bright-Glo luciferase kit (Promega), using a Glomax plate reader (Promega). Measurements were performed in duplicate and used to calculate 50% inhibitory concentrations (IC_50_) in GraphPad Prism software.

### Semi-quantitative ELISA

As described previously (O’Nions et al., 2020) nine columns of a half-well 96-well MaxiSorp plate were coated with purified SARS-CoV-2 Spike S1 protein in PBS (3 µg/ml per well in 25 µL) and the remaining three columns were coated with 25 μL goat anti-human F(ab)’2 diluted 1:1000 in PBS to generate the internal standard curve. After incubation at 4°C overnight, the ELISA plate was blocked for 1 hour in assay buffer (PBS, 5% milk, 0.05% Tween 20). Sera was diluted in assay buffer at dilutions from 1:50 to 1:5000 and 25 µL added to the ELISA plate. Serial dilutions of known concentrations of IgG standards were applied to the three standard curve columns in place of sera. The ELISA plate was then incubated for 2 hours at room temperature and then washed 4 times with PBS-T (PBS, 05% Tween 20). Alkaline phosphatase-conjugated goat anti-human IgG at a 1:1000 dilution was then added to each well and incubated for 1 hour. Following this, plates were washed 6 times with PBS-T and 25 µL of colorimetric alkaline phosphatase substrate added. Absorbance was measured at 405 nm. Antigen-specific IgG concentrations in serum were then calculated based on interpolation from the IgG standard results using a four-parameter logistic (4PL) regression curve fitting model.

### Serum samples

SAFER study samples were collected as previously described (Houlihan et al., 2020). Data derives from samples from 81 seropositive individuals during the first four months since infection. The study protocol was approved by the NHS Health Research Authority (ref 20/SC/0147) on 26 March 2020. Ethical oversight was provided by the South-Central Berkshire Research Ethics Committee. Serum samples from hospitalised patients were obtained during their hospital stay and through the Tissue Access for Patient Benefit (TAPb) scheme at The Royal Free Hospital (approved by UCL–Royal Free Hospital BioBank Ethical Review Committee Reference number: NC2020.24 NRES EC number: 16/WA/0289).

